# The Testing of Copper Intrauterine Devices to Mitigate Uterine Infections in Mares: A Preliminary Study

**DOI:** 10.1101/2024.11.01.621471

**Authors:** Carlos Gradil, Amelia Jennette, Klaus Becker, Teresa Haire, Carolynne Joone, Miranda Leibstein, Lisa Minter

## Abstract

This preliminary study evaluated the effectiveness of a novel hormone- and drug-free copper intrauterine device POD (CuiUPOD) with one to three copper-coated magnetic units in controlling *Streptococcus equi subsp. zooepidemicus* in seven intrauterine-inoculated mares. Initially, one unit was inserted into each mare and animals were followed weekly with trans-rectal ultrasonography, blood progesterone assay, uterine swabs for cytology and microbial culture, uterine specimens for biopsy, and trans-abdominal detection of the CuiUPOD using a handheld magnetic detector or a cell phone compass. Infection persisted after a CuiUPOD with one magnetic unit was inserted, but subsided shortly after two additional magnetic units were added. By day 60, at device removal, and after a 30-day exposure to copper, none of the mares presented with clinical signs of infection. On a scale of I-III (I, IIA, IIB, III), with I representing a healthy uterus, endometrial biopsies at the time of CuiUPOD removal, showed improved biopsy scores in two of seven mares (29%; *p* < 0.05). The *in vivo* experiments, complemented by an *in vitro* experiment, demonstrated a time- (24-96 hours) and dose-dependent response to Cu: three CuiUPODs - OD readings = 0.538; two CuiUPODs - OD = 0.513; and one CuiUPOD - OD = 0.452. As the concentration of copper increased, so did the antimicrobial effect. These findings suggest a promising use for the one-time application of a CuiUPOD with two or three magnetic units to mitigate uterine infections in mares.

**Simple Summary:** Only a limited number of antimicrobials are effective against most resistant bacteria. Several studies have demonstrated the biocidal effect of copper on bacteria, fungi, and viruses. Copper-containing intrauterine devices provide a non-pharmacological solution to prevent and treat uterine infections in equids. This preliminary study provides evidence of the antimicrobial properties of copper both in vivo and in vitro and explores its possible application in mare reproductive practice. A 30-day intrauterine exposure to copper showed no clinical signs of infection with a common bacterium in horses - *Streptococcus equi subsp. zooepidemicus* - in intrauterine-inoculated mares. As the concentration of copper increased, so did the antimicrobial effect. Copper intrauterine devices show promise in mitigating uterine infections in mares.

## 1. Introduction

The rise in antimicrobial resistance (AMR) is a concern both in veterinary and human medical fields and is one of the most significant indicators of universal One Health care.

New antibiotics are difficult to develop and would provide only temporary respite before resistance ensues. AMR infections are challenging to treat and create the need to develop drug-free treatment and prevention strategies. This is especially apparent within veterinary obstetrics and gynecology, due to the mare genital tract being prone to infections caused by more than one microbe. In certain circumstances, the “normal” microflora of the vagina and cervix can migrate into the uterus and cause an infection (review by Morris et al. 2020).

Egyptians first noted the antimicrobial effects of copper in 2600 BC (Dollwet, 1985). Copper is an effective biocidal surface material (Kuhn, 1983). Subsequent studies by many investigators have demonstrated the biocidal effect of copper on bacteria, fungi, and viruses (reviewed by Borkow and Gabbay, 2005, 2009). The effectiveness of copper extends to methicillin-resistant *Staphylococcus aureus* (Noyce, 2006; World Health Organization, 2010). Copper interacts with microorganisms causing cell death via membrane lipid peroxidation and degradation of nucleic acids (Erdogan and Rao, 2015).

Copper generates significant amounts of hydrogen peroxide and hydroxyl radicals that are very efficient biocides (Gunther et al. 1995).

Copper-containing intrauterine devices (IUDs) can be used in women for 10-12 years without adverse reactions (Goodwin et al. 2010). On the contrary, they effectively controlled yeast infections (Mishell 2010). Copper in uterine fluids of patients who used the copper T-380A intrauterine device from 6 months up to 3 years showed that total copper concentrations were 3.9-19.1 micrograms/ml. The copper leaching is in the form of complexes, avoiding the toxicity of copper in its free form (Timonen, 1976).

Even though the dissolution of the metal occurs via Cu(I) as Cu2O, it is oxidized to Cu(II) (Arancibia, et al. 2003).

Yeast infections account for 1–5% of all cases of endometritis in mares (reviewed by Scott, 2020) and are challenging to treat (Beltaire et al. 2012). CuiUPODs may address this challenge because copper ions leach from the CuiUPODs into the uterus. In preliminary experiments (unpublished data), copper quantitation by Mass Spectrometry in endometrial cytobrush samples from mares with CuiUPODs ranged from 0.16 to 0.198 µg/mL.

In this study, we explored the potential anti-microbial benefits of placing intra-uterine copper-containing pods in mares. We infused the most common bacterial pathogen responsible for endometritis in mares, *Streptococcus equi subsp. zooepidemicus (S. zooepidemicus*; Rasmussen et al., 2013) into the uterus and determined the efficacy of the CuiUPODs to treat or prevent uterine infection.

## 2. Materials and Methods

### 2.1. Animals

Six mares of various breeds, clinically normal, and 1 positive for *E. coli* were selected from the University herd for this 90-day study.

Mares were 14 ± 6.5 years (mean ± SD). Two mares were maiden and five were pluriparous. Experimental procedures were approved by the Institutional Animal Care and Use Committee protocol # 2262, University of Massachusetts Amherst. The authors declare that they have adhered to the Principles of Veterinary Medical Ethics of the American Veterinary Medical Association.

### 2.2. Intrauterine Device

An iUPOD is an intrauterine device made of 3 magnetic units (Fig. 1). The units can be left uncoated (Gradil et al. 2021) or coated with hormones, or antimicrobial or biocide chemical elements. The magnetic units can be used singly or in combination (1-3).

**Figure 1.**
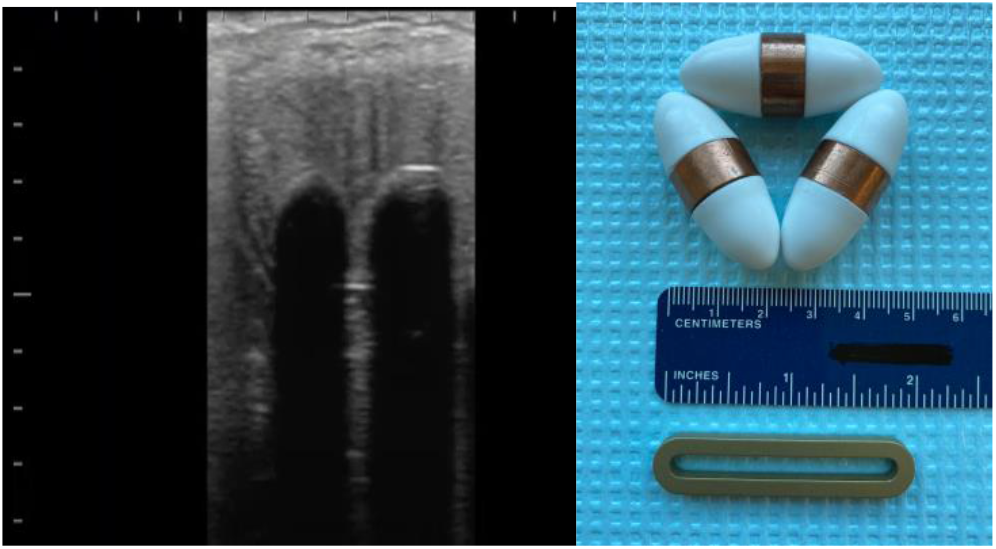
Left: Ultrasonic image of a CuiUPOD in the uterus. The left scale is in increments of 10 mm. Right: Copper CuiUPOD and the magnetic retriever. The size of the device is 40×16 mm with a mean unit weight of 26 g.

### 2.3. Experimental Design

A three-month experimental study with 1 to 3 copper-coated magnetic units CuiUPOD included the assessment of i) uterus and ovaries; ii) blood progesterone levels; iii) uterine cytology; iv) uterine culture; v) uterine health following device removal; and vi) trans-abdominal detection of the CuiUPOD, using a metal detector or a cell phone compass. The timelines of sampling for endometrial cytology, *S. zooepidemicus* inoculation, placement of the devices, uterine culture, and uterine biopsy, are shown in Tables 1, 2, and 3.

**Table 1.** Endometrial cytology Degrees of Inflammation.

|  | Pre-*S. zooepidemicus*<br>inoculation | *S. zooepidemicus* inoculation | CiUPOD treatment |  |
| --- | --- | --- | --- | --- |
| Mare # | Day -30 | Day 0 | Day 30<br>CiUPOD<br>Insertion | Day 60<br>CiUPOD<br>removal |
| 1 | None | High | High | Moderate |
| 2 | None | None | None | None |
| 3 | None | None | High | None |
| 4 | None | None | None | None |
| 5 | Moderate | Moderate | None | None |
| 6 | None | None | None | None |
| 7 | None | Moderate | None | None |

Inflammation was graded based on the number of polymorphonuclear neutrophils (PMNs) per five high-powered fields (240X). No Inflammation: <1 PMN/5 HPF; Moderate inflammation: 1-25 PMN/5 HPF; High inflammation: >200 PMN/5 HPF.

**Table 2.** Culture of endometrial swab samples.

|  | Pre-*S. zooepidemicus* inoculation | *S. zooepidemicus* inoculation | CiUPOD treatment |  |
| --- | --- | --- | --- | --- |
| Mare # | Day -30 | Day 0 | Day 30<br>CiUPOD<br>Insertion | Day 60<br>CiUPOD<br>removal |
| 1 | Negative | Positive<br>*5+*S. zooepidemicus* | Positive<br>5+*S. zooepidemicus* | Positive**<br>1+*S. zooepidemicus* |
| 2 | Negative | Negative | Negative | Negative |
| 3 | Negative | Negative | Positive<br>2+*S. zooepidemicus* | Negative |
| 4 | Negative | Negative | Negative | Negative |
| 5 | Positive<br>2+ *E. coli* | Positive<br>1+ *E. coli* | Negative | Negative |
| 6 | Negative | Negative | Negative | Negative |
| 7 | Negative | Negative | Negative | Negative |

\*Mare 1 tested positive for *S. zooepidemicus* (natural) infection before *S. zooepidemicus* inoculation, and was not inoculated subsequently. \*\*1+ Bacterial growth when compared to day 30 culture 5+ bacterial growth. \*≤ 1+ Considered clinically not significant.

**Table 3.** Horse endometrial biopsy.

| Mare ID | Day -7<br>(*S. zooepidemicus*<br>inoculation = Day 0) | Day 60<br>(iUPOD removal) |
| --- | --- | --- |
| 1 | IIA* | I |
| 2 | IIB | IIB |
| 3 | IIA | IIA |
| 4 | I | I |
| 5 | IIB* | IIA |
| 6 | I | I |
| 7 | IIA | IIA |

Changes in grade scores at the time of removal were as follows: i) increased - zero of seven (0%); decreased\* - two of seven (29%; $p > 0.05$ ); no change - five of seven (71%). Supplementary material link:
https://docs.google.com/document/d/1pnA4IdNKqAmCRaIcqVsY7-IH4II3Cjd7/edit?usp=share_link&oid=105117086797107510078&rtpof=true&sd=true

Uterine bacterial inoculation was used to determine if the inclusion of copper could treat or prevent uterine infections. The interval between bacterial inoculation and CuiUPOD treatment was 30 days. This interval provided ample opportunity for mares to cycle post-bacterial inoculation, and to determine if they could clear the bacteria independently. The inoculum used in the *in vivo* and *in vitro* experiments was a genital strain kindly provided by Prof. Erdal Erol, Veterinary Diagnostic Laboratory, University of KY.

Mares were examined before the experiment by routine ultrasonography for pregnancy, presence of uterine fluid, urine pooling, granulosa cell tumor, etc. The perineal area was scrubbed with 7.5% povidone-iodine for guarded endometrial swabbing for endometrial cytology (Joone CJ et al. 2021), microbial culture, uterine specimens for biopsy, CuiUPOD insertion or retrieval. Mares were sedated with 0.02-0.04 mg/kg detomidine hydrochloride^1a^, administered intravenously.

Sampling for culture was retrieved on day -30, before intrauterine inoculation with an equine strain of 100,000 colony-forming units of *S. zooepidemicus* in 10 mL sterile saline on days 0, 30, and 60.

Clinical signs of infection, for example, the presence of intrauterine fluid (IUF) in a mare in diestrus (McCue PM, 2008) were monitored by ultrasonography^1b^ and confirmed by uterine culture. To determine uterine inflammation and infection, uterine swabs for cytology and bacterial culture were retrieved every 30 days. To determine uterine inflammation and integrity caused by the devices, paired uterine specimens for biopsy were retrieved on days -7 and 60. On day 30, the CuiUPODs were inserted trans-cervically into seven mares at random stages of the estrous cycle. All CuiUPOD units were removed on day 60.

### 2.4. Monitoring the Estrous Cycle

Mares were monitored weekly by ultrasonography and laboratory analysis of progesterone values (Aurich C, 2011). The number and size of follicles, the corpora lutea, and the presence of IUF were assessed with ultrasonography and recorded. Dorsoventral depth measurements were used to determine the amount of IUF.

### 2.5. Laboratory Analysis of Blood, Cytology, Bacterial Culture, and Biopsy

Serum samples for progesterone (P4) analysis were collected on day -30, and then weekly for 12 weeks and sent to a reference laboratory^c^. Samples were analyzed using MP Biomedicals Progesterone RIA, without previous knowledge of treatment status. Reference progesterone values were as follows: Absence of active luteal tissue: 0.1–0.5 ng/mL - Estrus; borderline for the presence of luteal tissue: 0.5–1.0 ng/mL; the presence of luteal tissue: > 1.0 ng/mL - Diestrus.

Sterilized double-guarded endometrial swabs^d^ were obtained for endometrial cytology and culture from all mares on days -30, 0, 30, and 60 - *immediately before* CuiUPOD removal. Diff-Quick^®e^ stain was used for cytological assessment. Endometrial swabs were streaked on a blood agar plate and incubated for 48 hours at 37° C. Dispersed pure colony growth of *S. zooepidemicus* was observed in the absence of hemolysis^c^.

Paired biopsy samples were collected on day -7, and *immediately before* CuiUPOD removal on day 60, and evaluated by a board-certified veterinary pathologist, blinded to treatment history, and graded as described by Kenney and Doig (1986).

### 2.6. Monitoring Device Retention

Mares were examined by trans-rectal ultrasonography within five minutes post-device insertion, as shown in Fig. 1.

CuiUPODs were detected using a magnetic detector^2f^ or a cell phone compass on day 30 and weekly thereafter. The reliability of the metal detector was 100%, i.e., true alarm alerts (Gradil et al. 2021), and likewise for the cell phone compass.

### 2.7. In vitro Bacterial Growth Response to CiUPODs

Eight samples were created as follows: 1. Muller Hinton (MH) broth (Millipore, Burlington, MA, Cat. Nr. 90922) supplemented with 1% horse serum and enriched with 0.5g/100mL Hematin (Millipore, Burlington, MA, Cat. H3281) for selective growth support (Ritchey and Seeley, 1976) with NO *S. zooepidemicus* or devices; 2. MH broth with 1% horse serum with *S. zooepidemicus* and NO devices; 3. MH broth with 1% horse serum with *S. zooepidemicus* and 1 CiUPOD; 4. MH broth with 1% horse serum with *S. zooepidemicus* and 2 CiUPODs; 5. MH broth with 1% horse serum with *S. zooepidemicus a*nd 3 CiUPODs; 6. MH broth with 1% horse serum with 3 Teflon IUD; 7. MH broth with 1% horse serum NO *S. zooepidemicus* and 3 CiUPODs; 8. MH broth with 1% horse serum NO *S. zooepidemicus* and 3 Teflon devices.

Each sample was contained in an aseptic culture container. MH broth with 1% horse serum was created and filtered to ensure sterility. Devices were immersed in 70% isopropanol for 1 hour and allowed to dry before the experiments. Thirty-six mL of MH broth was added to each cup to ensure total immersion of the devices. Samples were then placed on a shaker at 120 bpm to mimic a working trot/working canter at 37°C to mimic the average core temperature of the horse. Photos were taken at timepoints 24, 48, 72, and 96 hours to observe any physical changes between the samples. A Gram stain was performed at 96 hours to confirm that it was a gram-positive bacterium.

Additionally, ocular density measurements were performed to determine the overall concentration (Maia et al., 2016) of the *S. zooepidemicus* at 620 nm utilizing a Biotek, Synergy 2^f^, instrument reader with turbidity measurement capabilities using a standard blank broth-only.

## 3. Results

### 3.1 Monitoring the Estrous Cycle and IUF in the Uterine Lumen

Trans-rectal ultrasonography was conducted on day -30, and again, weekly. CLs were observed in both ovaries of all mares. Mares #1, 2, and 3 double-ovulated during one estrous cycle. The average diestrus length in days for seven mares was (13.2 ± 4.7), the average estrus was (8.9 ± 6.4), and the interestrus interval was (22.1 ± 7.4). Mares # 4, 6, and 7 had extended periods of diestrus (16, 21, and 28 days, respectively) following CuiUPOD insertion.

Mare 1 contracted an *S. zooepidemicus* natural infection and was not inoculated. Two mares had IUF: mare 1 (days 8, 15, 17); and mare 3 (days 1, 3, 4, 10, 30). There was no IUF in six of seven mares at CuiUPOD, except in mare 3 with non-echogenic fluid near the device and negative culture. IUF was absent in all the mares six weeks post-CuiUPOD removal.

### 3.2. Uterine Cytology

### 3.3. Uterine Culture

The results of endometrial swabs collected on days -30, 0, 30, and 60 are presented in Table

### 3.4 Estrous Cycle Periodicity - Progesterone Patterns

On day 0, three of seven mares were in estrus (P4 < 1 ng/mL). On day 30 (CuiUPOD insertion), there were two of seven mares with (P4 < 1 ng/mL) and five of seven mares with P4 > 1 ng/mL. Three of seven mares fitted with the device had an active CL for periods >14 days with P4 > 1 ng/mL, and in one of seven mares, there were periods >14 days with P4 concentrations of < 1 ng/mL.

### 3.5. Uterine Health following Device Removal

Biopsy results are shown in Table 3.

### 3.6. Device Retention

The magnetic units self-aggregate in the uterus, as illustrated in Fig. 1.

Mare 4 (nulliparous) lost a single CuiUPOD magnetic unit on day 37. Two replacement CuiUPOD magnetic units were reinserted immediately to ascertain if two units would be retained through day 60.

### 3.7. In vitro Bacterial Growth Response to CiUPODs

We observed a time (24-96 hours) and dose-dependent bacterial growth response to Cu: three CuiUPODs - OD readings = 0.538; two CuiUPODs - OD = 0.513; and one CuiUPOD - OD = 0.452 Fig. 2.

**Figure 2.**
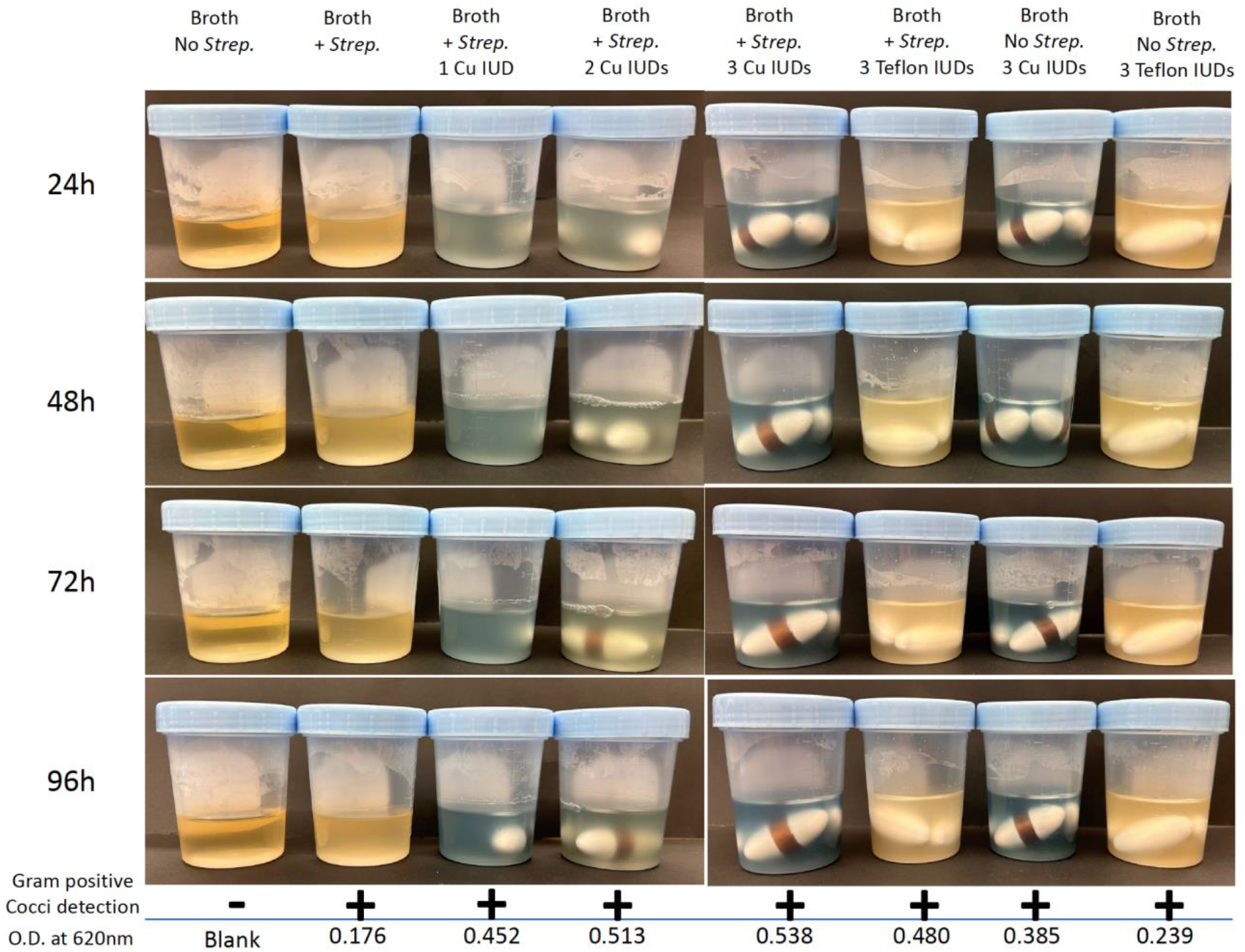
Gallery of all the samples and time points, with the final optical density (OD) reading at time point 96 hours. OD decreases as bacterial concentration decreases, due to exposure to copper meaning more bacteria lead to a higher OD reading because of copper in the PODs.

## 4. Discussion

The reduced bacterial growth and absence of clinical signs of infection after the insertion of CuiUPODs in mares 1 and 3 suggest that CuiUPODs may successfully treat both experimental and natural intrauterine infection, as *S. zooepidemicus* that is common in endometritis, was present in mare 1. Infection persisted after a CuiUPOD with one magnetic unit was inserted but subsided shortly after two additional magnetic units were administered to mares. This suggests a higher copper concentration is more effective in treating established infections.

In the current study, three of seven mares fitted with the device had an active CL, as determined by progesterone assay, for periods >14 days. In previous studies on estrus suppression (Katila T, 2015); Nie et al., 2003), the prolongation of CL function was one month longer with glass balls than plastic balls. Future studies should compare the effect of copper with non-copper devices on the estrous cycle.

At the time of CuiUPOD removal, there was an improvement in biopsy scores in two of seven mares (29%; p >0.05), which supports previous findings (Gradil et al. 2019).

This research was limited in duration and sample size, the heterogeneous selection of mares, and the need for more fertility data post-treatment. The study was carried out on mares inoculated with *S. zooepidemicus*, rather than with mares positively diagnosed with bacterial or fungal endometritis. Mare #5, the oldest multiparous mare in the study, and despite poor perineal anatomy, still cleared the *E.coli* infection. Poor perineal anatomy decreases the ability of the vulva to act as a barrier against ascending infection. It may facilitate aspiration of air into the vagina with increased predisposition to uterine infections and reinfection (McCue, 2008). In this case, the mare remained clear of infection for the latter half of the study. Whether this was due to the presence of the device is difficult to ascertain.

Future studies should include higher concentrations of copper. This can be achieved by entirely cladding or encapsulating each unit, which increases the surface area per unit by 75.6 %. This is a significant increase in Cu^2+^ availability with the potential to decrease exposure and treatment time.

## 5. Conclusion

This preliminary study provides evidence of the antimicrobial properties of copper both *in vivo* and *in vitro* and explores its possible application in mare reproductive practice.

A 30-day in vivo exposure to copper resulted in no significant clinical signs of *Streptococcus equi subsp. zooepidemicus* infection in intrauterine-inoculated mares. The reduced bacterial growth in mare 1, when compared to day 30 culture (5+ bacterial growth) and the absence of clinical signs of infection (≤ 1+; clinically not significant) after the insertion of CuiUPODs, together with the negative culture in mare 3 suggest that CuiUPODs may successfully treat both experimental and natural intrauterine infection.

As the concentration of copper was increased in bacterial cultures, *in vitro*, so did the antimicrobial effect. Altogether our data show that CuiUPODs hold promise in mitigating uterine infections in mares.

## Acknowledgments

The authors would like to thank A. Ruben, A. Golembeski, and the University of Massachusetts Amherst Hadley Farm staff for their help with this project, and C. Morcom for providing access to mares. The prototype device was kindly provided by Amazing Magnets, TX.

## Author Contributions

Carlos Gradil: Conceptualization, Formal analysis, Writing, Visualization, Project administration, Funding acquisition. Amelia Jennett: Sample collection, Bacterial culture for in vivo studies, Visualization. Klaus Becker: Methodology, Bacterial culture, and analysis. Carolynne Joone: Review and editing. Teresa Haire: Project administration and supervision. Miranda Leibstein: In vitro studies, Visualization. Lisa Minter: Review and editing. All authors have read and agreed to the published version of the manuscript.

## Funding

This work was supported by revenue from UMass Equine Reproductive Services and MAS00513, University of Massachusetts Amherst.

## Data availability

The authors declare that data supporting the findings of this study are available within the article.

## Ethical approval

All experimental procedures were approved by the University of Massachusetts Amherst Institutional Animal Care and Use Committee; all work adhered to the Principles of Veterinary Medical Ethics of the American Veterinary Medical Association.

## Competing interests

The authors declare no competing interests.

---

**Table 1.** Endometrial Cytology.

|  | Pre-<u>S.</u><br><u>zooepidemicus</u><br><u>inoculation</u> |  | Post-treatment |  |
| --- | --- | --- | --- | --- |
| Mare # | Day 0 | Day 30 | Day 60<br>CiUPOD<br>Insertion | Day 90<br>CiUPOD<br>Retrieval |
| 1 | < than 1 PMN<br>per 5 high<br>powered fields | 25 - 200 PMN per 5<br>high powered fields | 25 - 200 PMN per 5<br>high powered fields | < than 25 PMN per 5<br>high powered fields |
| 2 | < than 1 PMN<br>per 5 high<br>powered fields | < than 1 PMN per 5<br>high powered fields | < than 1 PMN per 5<br>high powered fields | < than 1 PMN per 5<br>high powered fields |
| 3 | < than 1 PMN<br>per 5 high<br>powered fields | < than 1 PMN per 5<br>high powered fields | 25 - 200 PMN per 5<br>high powered fields | < than 1 PMN per 5<br>high powered fields |
| 4 | < than 1 PMN<br>per 5 high<br>powered fields | < than 1 PMN per 5<br>high powered fields | < than 1 PMN per 5<br>high powered fields | < than 1 PMN per 5<br>high powered fields |
| 5 | < than 25 PMN<br>per 5 high<br>powered fields | < than 25 PMN per 5<br>high powered fields | < than 1 PMN per 25<br>high powered fields | < than 1 PMN per 5<br>high powered fields |
| 6 | < than 1 PMN<br>per 5 high<br>powered fields | < than 1 PMN per 5<br>high powered fields | < than 1 PMN per 5<br>high powered fields | < than 1 PMN per 5<br>high powered fields |
| 7 | < than 1 PMN<br>per 5 high<br>powered fields | < than 25 PMN per 5<br>high powered fields | < than 1 PMN per 5<br>high powered fields | < than 1 PMN per 5<br>high powered fields |

On day 30 three mares had no signs of inflammation and 4 of seven mares had few numbers of polymorphonuclear neutrophils (PMNs.) On day 60, the time of iUPOD insertion, three mares had no signs of inflammation, two mares had few numbers of PMNs, and two mares had moderate signs of inflammation. On day 90, the day of iUPOD retrieval, six mares had no signs of inflammation and 1 mare had mild signs of inflammation. The inflammatory cells consisted primarily of mature hyper-segmented PMNs.

**Table 2.** Culture of endometrial swab samples.

| Mare # | Day 0 | Day 30 | Day 60 | Day 90 |
| --- | --- | --- | --- | --- |
| 1 | Negative | Positive<br>**5<sup>+</sup> <u>S. zooepidemicus</u> | Positive<br>5<sup>+</sup> <u>S. zooepidemicus</u> | Positive*<br>1<sup>+</sup> <u>S. zooepidemicus</u> |
| 2 | Negative | Negative | Negative | Negative |
| 3 | Negative | Negative | Positive<br>2<sup>+</sup> <u>S. zooepidemicus</u> | Negative |
| 4 | Negative | Negative | Negative | Negative |
| 5 | Positive<br>2<sup>+</sup> <u>E. coli</u> | Positive<br>1<sup>+</sup> <u>E. coli</u> | Negative | Negative |
| 6 | Negative | Negative | Negative | Negative |
| 7 | Negative | Negative | Negative | Negative |

**Table 3.** Horse Endometrial Biopsy.

| Day -7 | Mare # | Inflammation |
| --- | --- | --- |
| TDP21-130-3 | 1 | Inflammation ranges from minimal to moderate, multifocal, predominantly lymphoplasmacytic, |
|  | 2 | Mild to moderate, multifocal, chronic (lymphoplasmacytic) inflammation with numerous hemosiderin-laden macrophages |
|  | 3 | Mild, diffuse lymphoplasmacytic inflammation |
|  | 4 | Minimal to mild, diffuse lymphoplasmacytic inflammation |
|  | 5 | Mild to moderate, multifocal, chronic (lymphoplasmacytic) inflammation |
|  | 6 | Minimal |
|  | 7 | Mild, diffuse lymphoplasmacytic inflammation |
| **Day 60** |  |  |
| TDP21-130-4 | 1 | Minimal, scattered, multifocal, chronic lymphoplasmacytic |
|  | 2 | Mild to moderate, multifocal, chronic (lymphoplasmacytic) inflammation with numerous hemosiderin-laden macrophages |
|  | 3 | Mild to moderate, multifocal, chronic lymphoplasmacytic |
|  | 4 | Minimal to mild, diffuse lymphoplasmacytic inflammation |
|  | 5 | Mild to moderate, multifocal, chronic lymphoplasmacytic |
|  | 6 | Minimal, scattered, multifocal, chronic lymphoplasmacytic |
|  | 7 | Mild to moderate, multifocal, chronic lymphoplasmacytic |

| Day -7 | Mare # | Fibrosis |
| --- | --- | --- |
| TDP21-130-3 | 1 | Mild to moderate fibrosis, with rare fibrotic glandular nests, and significant variability in location |
|  | 2 | Moderate to marked, diffuse fibrosis with multifocal fibrotic glandular nesting |
|  | 3 | Mild to moderate fibrosis with frequent fibrotic glandular nests throughout the sample |
|  | 4 | Mild fibrosis, no overt nesting |
|  | 5 | Mild to moderate fibrosis, with +++ fibrotic glandular nests (>5 per 5mm linear field) |
|  | 6 | NA |
|  | 7 | Mild to moderate fibrosis with frequent fibrotic glandular nests throughout the sample |
| **Day 60** |  |  |
| TDP21-130-4 | 1 | Minimal and scattered |
|  | 2 | Moderate to marked, diffuse fibrosis with multifocal fibrotic glandular nesting |
|  | 3 | Mild to moderate fibrosis with fibrotic glandular nesting |
|  | 4 | Mild fibrosis, no overt nesting |
|  | 5 | Mild to moderate fibrosis with fibrotic glandular nesting |
|  | 6 | Minimal fibrosis, with a single focus of nesting |
|  | 7 | Mild to moderate fibrosis with fibrotic glandular nesting |

| Day -7 | Mare # | Score: | Comments |
| --- | --- | --- | --- |
| TDP21-130-3 | 1 | IIA | There is variability of the lesions multifocally throughout the sample, with areas with intense inflammation and fibrosis, and areas with no significant changes |
|  | 2 | IIB | Many hemosiderin-laden macrophages (siderophages), suggest chronic hemorrhage/inflammation or postpartum uterus. Changes in the vascular adventitia consistent with elastosis (possibly age related) |
|  | 3 | IIA |  |
|  | 4 | I |  |
|  | 5 | IIB* | The amount of fibrosis would push this sample to III, however, inflammation is not severe |
|  | 6 | I | Multifocal hemorrhage, likely artifactual |
|  | 7 | IIA | Mild, multifocal endometrial glandular atrophy |
| **Day 60** |  |  |  |
| TDP21-130-4 | 1 | I |  |
|  | 2 | IIB | Many hemosiderin-laden macrophages (siderophages), suggest chronic hemorrhage/inflammation or postpartum uterus. Changes in the vascular adventitia consistent with elastosis (possibly age related) |
|  | 3 | IIA |  |
|  | 4 | I |  |
|  | 5 | IIA |  |
|  | 6 | I | Nesting is in a single focus, and due to minimal inflammation |
|  | 7 | IIA |  |

## Footnotes

1 a. Dormosedan®, Zoetis, NJ 07054-3825. b. Hitachi Aloka Medical America Inc, CT 06492-5903. c. Cornell University Animal Health Diagnostic Centre, NY 14850. d. Breeders Choice, Irwindale, CA 91706. e. Diff-Quick® American Scientific Products, OH 43235. f. Agilent Technologies, Santa Clara, CA 95051.

2 f. PD240CB, CEIA USA Ltd, OH 44236.

